## Supplementary Fig. for "Variant antigen diversity in *Trypanosoma vivax* is not driven by recombination"

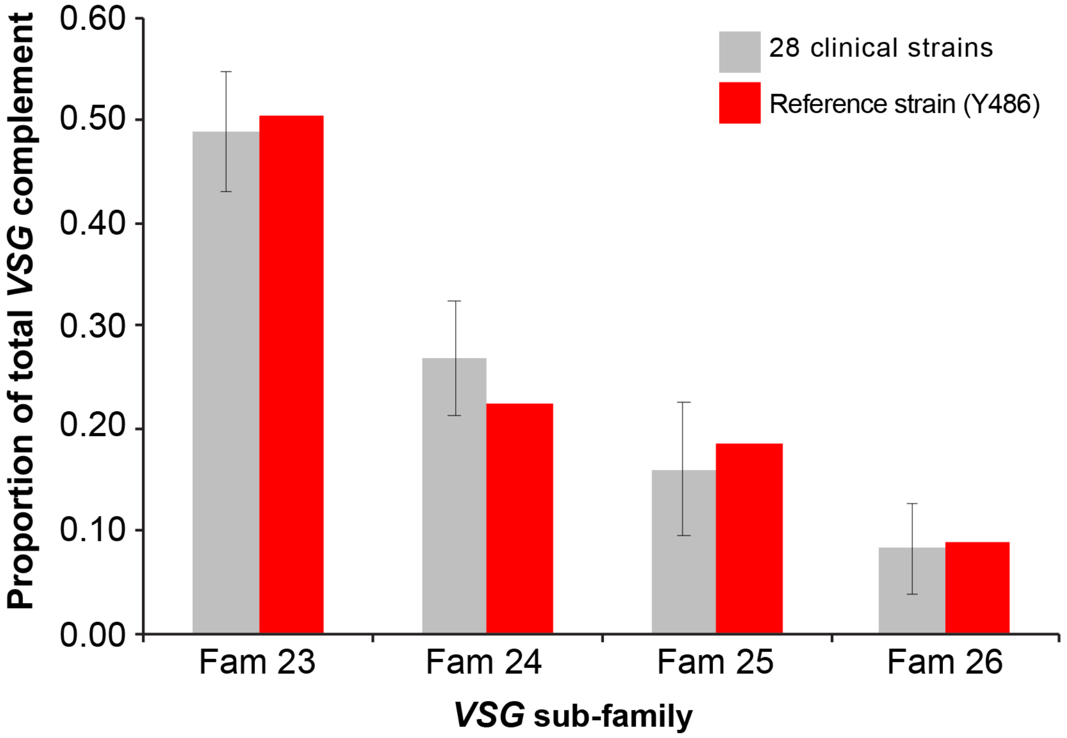


**Supplementary Fig. 1**. **Proportions of four conserved *VSG* sub-families (Fam23-26) to total *VSG* complement in 28 *T. vivax* clinical strains compared with the *T. vivax* Y486 reference strain.** Previously, we established Fam23-26 in *T. vivax* Y486[1]. These were observed in all clinical strain genome sequences and in approximately the same relative proportions. This makes the sub-families unsuitable for discriminating among strains and therefore as a basis for variant antigen profiling, and requires the use of more variable taxa (i.e. phylotypes).

# P24

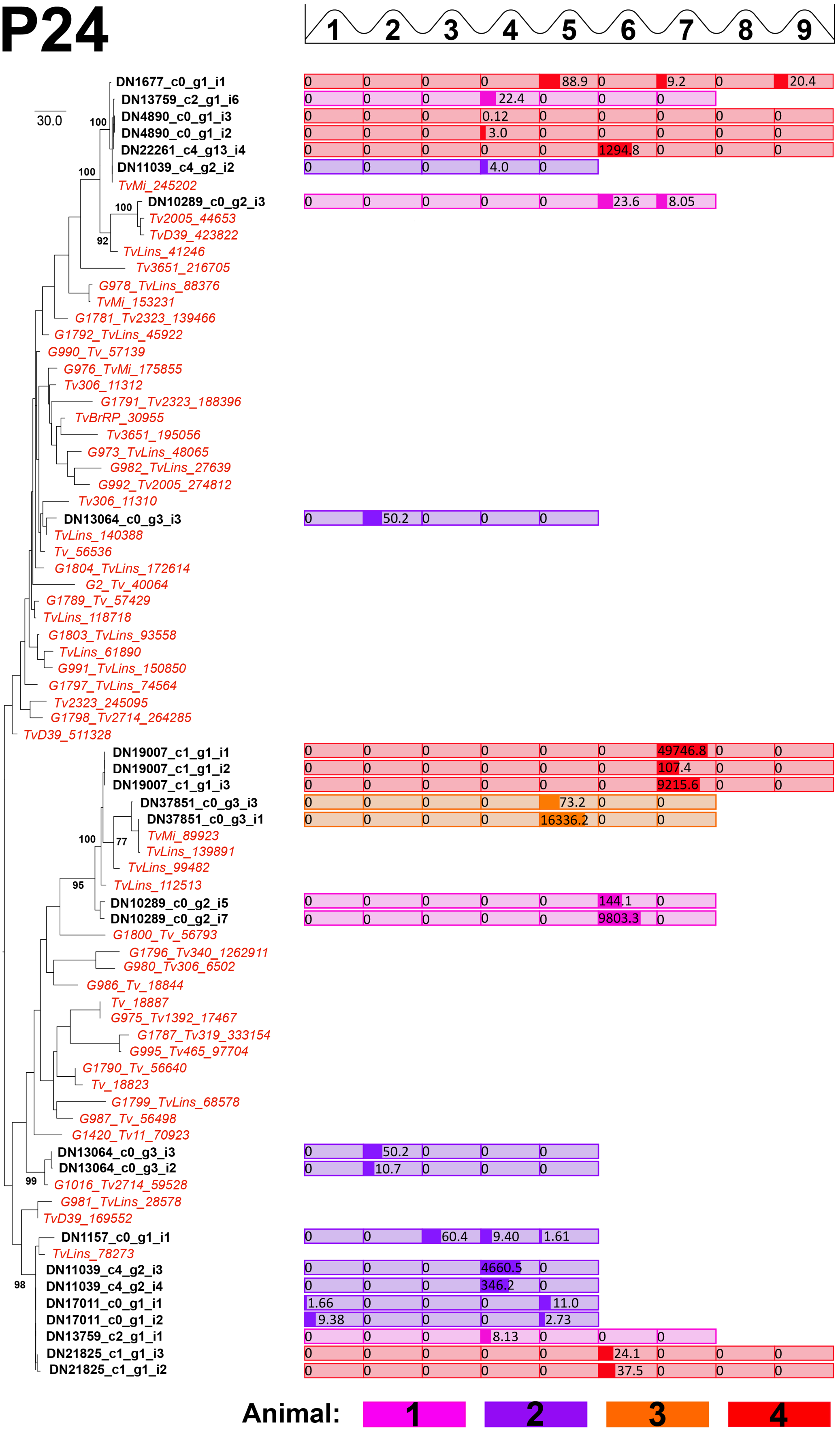

1 2 3 4 5 6 7 8 9
30.0
DN1677_c0_g1_i1 0 0 0 0 88.9 0 9.2 0 20.4
DN13759_c2_g1_i6 0 0 0 22.4 0 0 0
DN4890_c0_g1_i3 0 0 0 0.12 0 0 0 0 0
DN4890_c0_g1_i2 0 0 0 3.0 0 0 0 0 0
DN22261_c4_g13_i4 0 0 0 0 0 1294.8 0 0 0
DN11039_c4_g2_i2 0 0 0 4.0 0
TvMi_245202
DN10289_c0_g2_i3 0 0 0 0 0 23.6 8.05
Tv2005_44653
TvD39_423822
TvLins_41246
Tv3651_216705
G978_TvLins_88376
TvMi_153231
G1781_Tv2323_139466
G1792_TvLins_45922
G990_Tv_57139
G976_TvMi_175855
Tv306_11312
G1791_Tv2323_188396
TvBrRP_30955
Tv3651_195056
G973_TvLins_48065
G982_TvLins_27639
G992_Tv2005_274812
Tv306_11310
DN13064_c0_g3_i3 0 50.2 0 0 0
TvLins_140388
Tv_56536
G1804_TvLins_172614
G2_Tv_40064
G1789_Tv_57429
TvLins_118718
G1803_TvLins_93558
TvLins_61890
G991_TvLins_150850
G1797_TvLins_74564
Tv2323_245095
G1798_Tv2714_264285
TvD39_511328
DN19007_c1_g1_i1 0 0 0 0 0 0 49746.8 0 0
DN19007_c1_g1_i2 0 0 0 0 0 0 107.4 0 0
DN19007_c1_g1_i3 0 0 0 0 0 0 9215.6 0 0
DN37851_c0_g3_i3 0 0 0 0 73.2 0 0
DN37851_c0_g3_i1 0 0 0 0 16336.2 0 0
TvMi_89923
TvLins_139891
TvLins_99482
TvLins_112513
DN10289_c0_g2_i5 0 0 0 0 0 144.1 0
DN10289_c0_g2_i7 0 0 0 0 0 9803.3 0
G1800_Tv_56793
G1796_Tv340_1262911
G980_Tv306_6502
G986_Tv_18844
Tv_18887
G975_Tv1392_17467
G1787_Tv319_333154
G995_Tv465_97704
G1790_Tv_56640
Tv_18823
G1799_TvLins_68578
G987_Tv_56498
G1420_Tv11_70923
DN13064_c0_g3_i3 0 50.2 0 0 0
DN13064_c0_g3_i2 0 10.7 0 0 0
G1016_Tv2714_59528
G981_TvLins_28578
TvD39_169552
DN1157_c0_g1_i1 0 0 60.4 9.40 1.61
TvLins_78273
DN11039_c4_g2_i3 0 0 0 4660.5 0
DN11039_c4_g2_i4 0 0 0 346.2 0
DN17011_c0_g1_i1 1.66 0 0 0 11.0
DN17011_c0_g1_i2 9.38 0 0 0 2.73
DN13759_c2_g1_i1 0 0 0 8.13 0 0 0
DN21825_c1_g1_i3 0 0 0 0 0 24.1 0 0 0
DN21825_c1_g1_i2 0 0 0 0 0 37.5 0 0 0
100 100 100 92 100 77 95 99 98
Animal: 1 2 3 4

**Supplementary Fig. 2. Maximum likelihood phylogeny of Phylotype 24 showing the relationships among constituent genes from reference and strain genomes (COG type sequences, shown in red) and expressed *VSG* sequences from *T. vivax* Lins (N = 25; shown in black).** The tree was estimated from a 585 bp amino acid alignment using a GTR+$\Gamma$ model in RAXML[2]. Some gene sequences have been removed because they were too short. Robustness values (100 non-parametric bootstraps) are shown beside selected internal nodes. Transcript abundance values (CPM) for each peak of parasitaemia are shown beside their cognate terminal nodes. These values are shaded by animal replicate.

P24 was observed at 15/29 peaks across four replicates and comprised an average of 2.33±1.3 transcripts per observation. Phylotypes were routinely observed to comprise multiple, distinct transcripts. The maximum number of unique P24 transcripts observed at a single peak was five (A3, peak 4). P24 comprised a single transcript on 5/15 peaks when it was observed.

Individual phylotypes were reproducibly expressed in different animals, and P24 provides an example of this. P24 was unique among phylotypes by providing a superabundant, dominant *VSG* in all animals in late infection (see Figure 4; marked here with an asterisk). However, the individual transcripts implicated in different animals are not identical. They are sometimes very closely related (e.g. superabundant P24 transcripts in A1 (DN10289) and A4 (DN19007) share 94.9% nucleotide identity), but elsewhere they are more distinct (e.g. the superabundant P24 transcripts in A2 (DN11039) and A3 (DN37851) share only 73.5% nucleotide identity).

In most cases where a phylotype was observed in multiple animals, the transcripts concerned were not identical. However, the figure also provides examples of identical transcripts expressed in multiple animals, for example, transcript DN13759_c2_g1_i1 in A1 and DN11039_c4_g2_i4 in A2, as well as DN11039_c4_g2_i2 in A2 and DN22261 in A4.

Phylotypes were observed to persist across consecutive peaks in the same animal. P24 provides examples of this, e.g. it was observed at every peak in A2, being superabundant at peaks 4 and 5. Nine different transcripts contribute to this profile, and none of these persist throughout the experiment. However, the figure does provide examples of individual *VSG* persisting across peaks. For example, DN1677 was expressed at peaks 6, 8 and 10 in A4, DN1157 was expressed at peaks 4, 5 and 6 in A2, and DN10289_c0_g2_i3 was expressed at both peaks 6 and 7 in A1.

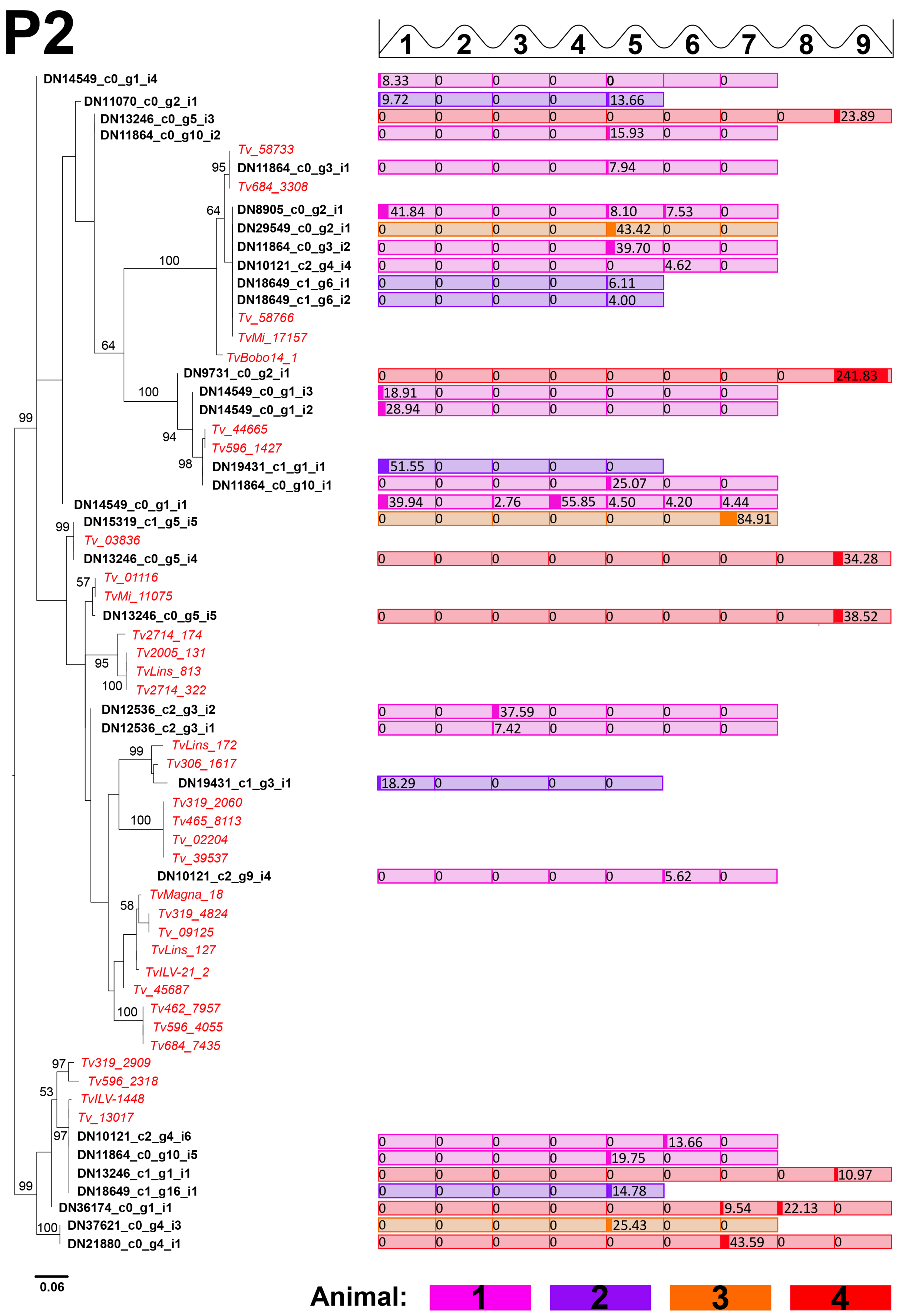
P2
1 2 3 4 5 6 7 8 9
DN14549_c0_g1_i4
DN11070_c0_g2_i1
DN13246_c0_g5_i3
DN11864_c0_g10_i2
Tv_58733
DN11864_c0_g3_i1
Tv684_3308
DN8905_c0_g2_i1
DN29549_c0_g2_i1
DN11864_c0_g3_i2
DN10121_c2_g4_i4
DN18649_c1_g6_i1
DN18649_c1_g6_i2
Tv_58766
TvMi_17157
TvBobo14_1
DN9731_c0_g2_i1
DN14549_c0_g1_i3
DN14549_c0_g1_i2
Tv_44665
Tv596_1427
DN19431_c1_g1_i1
DN11864_c0_g10_i1
DN14549_c0_g1_i1
DN15319_c1_g5_i5
Tv_03836
DN13246_c0_g5_i4
Tv_01116
TvMi_11075
DN13246_c0_g5_i5
Tv2714_174
Tv2005_131
TvLins_813
Tv2714_322
DN12536_c2_g3_i2
DN12536_c2_g3_i1
TvLins_172
Tv306_1617
DN19431_c1_g3_i1
Tv319_2060
Tv465_8113
Tv_02204
Tv_39537
DN10121_c2_g9_i4
TvMagna_18
Tv319_4824
Tv_09125
TvLins_127
TvILV-21_2
Tv_45687
Tv462_7957
Tv596_4055
Tv684_7435
Tv319_2909
Tv596_2318
TvILV-1448
Tv_13017
DN10121_c2_g4_i6
DN11864_c0_g10_i5
DN13246_c1_g1_i1
DN18649_c1_g16_i1
DN36174_c0_g1_i1
DN37621_c0_g4_i3
DN21880_c0_g4_i1
0.06
Animal: 1 2 3 4

**Supplementary Fig. 3. Maximum likelihood phylogeny of Phylotype 2 showing the relationships among constituent genes from reference and strain genomes (COG type sequences, shown in red) and expressed *VSG* sequences from *T. vivax* Lins (N = 31; shown in black).** The tree was estimated from a 228 bp alignment using a GTR+Γ model in RAXML[2]. Robustness values (100 non-parametric bootstraps) are shown beside selected internal nodes. Beside terminal nodes representing expressed *VSG* are the transcript abundance values (CPM) for each peak of parasitaemia. These values are shaded by animal replicate.

P2 was observed at 13/29 peaks across four replicates and comprised an average of 3.08±1.9 transcripts per observation. Phylotypes were routinely observed to comprise multiple, distinct transcripts. P2 comprised a single transcript on 4/13 peaks when it was observed. The maximum number of unique P2 transcripts observed at a single peak was seven (A2, peak 5).

Individual phylotypes were reproducibly expressed in different animals, and P2 provides an example of this, e.g. it is superabundant in both A1 and A2 at peak 1. However, this implicates five different P2 transcripts in A1, but a single transcript in A2 (DN19431_c1_g3_i1), which was only seen in that animal. While this is typical, the figure also provides examples of identical transcripts being expressed in different animals. For example, DN10121_c2_g4_i6 in A1 was identical to DN13246_c1_g1_i1 in A4 and DN18649_c1_g16_i1 in A2. Similarly, DN13246_c0_g5_i3 in A4 and DN11864_c0_g10_i2 in A4 are identical.

Phylotypes were observed to persist across consecutive peaks in the same animal. P2 provides a rare example in which the same transcript is expressed. DN14549_c0_g1_i1 was expressed throughout the experiment in A1 except at peak 2. While it was among the dominant VSG at peaks 1 and 4, it persisted at lower levels for the rest of the experiment, apparently being supplanted by other P2 transcripts in abundance. In other cases, individual transcripts re-emerged late in the experiment after early expression, e.g. DN8905 was the dominant *VSG* in A1 at peak 1, and re-emerged at moderate levels during peaks 5 and 6, while DN11070 was expressed in A2 at peak 2 and re-emerged at peak 6 (as a low abundance transcript in both cases).

# P40

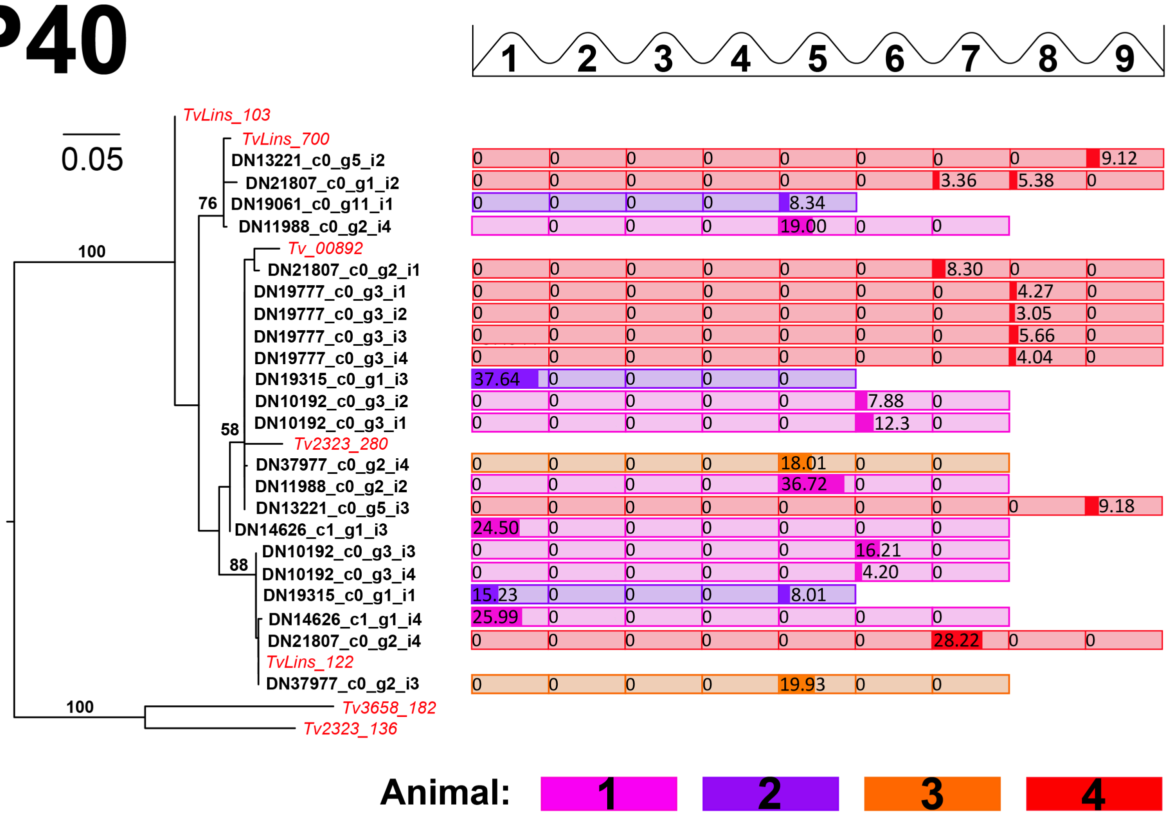


**Supplementary Fig. 4. Maximum likelihood phylogeny of Phylotype 40 showing the relationships among constituent genes from reference and strain genomes (COG type sequences, shown in red) and expressed *VSG* sequences from *T. vivax* Lins (shown in black).** The tree was estimated from a 489 bp alignment using a GTR+Γ model in RAXML[2]. Robustness values (100 non-parametric bootstraps) are shown beside selected internal nodes. Beside terminal nodes representing expressed *VSG* are the transcript abundance values (CPM) for each peak of parasitaemia. These values are shaded by replicate animal.

P40 was observed at 9/29 peaks across four replicates and comprised an average of 2.67±1.1 transcripts per observation. Phylotypes were routinely observed to comprise multiple, distinct transcripts. The maximum number of unique P40 transcripts observed at a single peak was five (A4, peak 9). P40 comprised a single transcript on 2/9 peaks when it was observed.

Individual phylotypes were reproducibly expressed in different animals, and P40 provides an example of this, e.g. it is superabundant in both A1 and A2 at peak 1 (different transcripts), and co-dominant at peak 5 in A3 and A1. Again, the actual transcripts in the two animals are different. However, there are several examples of identical transcripts being observed as low abundance forms in multiple animals, e.g. DN19777 in A4 and DN10192 in A1 are identical to the superabundant DN19315_c0_g1_i3 in A2.

Phylotypes were observed to persist across consecutive peaks in the same animal. P40 provides examples of this, e.g. across peaks 8-10 in A4, where it was superabundant at peak 8 and then present as multiple, low abundance transcripts thereafter. it was observed at every peak in A2, being superabundant at peaks 4 and 5. Typically when phylotypes persist or re-emerge, the actual transcripts concerned are distinct. However, transcript DN19315_c0_g1_i1 is expressed at peak 1 in A2, then again at peak 5, providing a rare example of the same transcript re-emerging during the experiment.

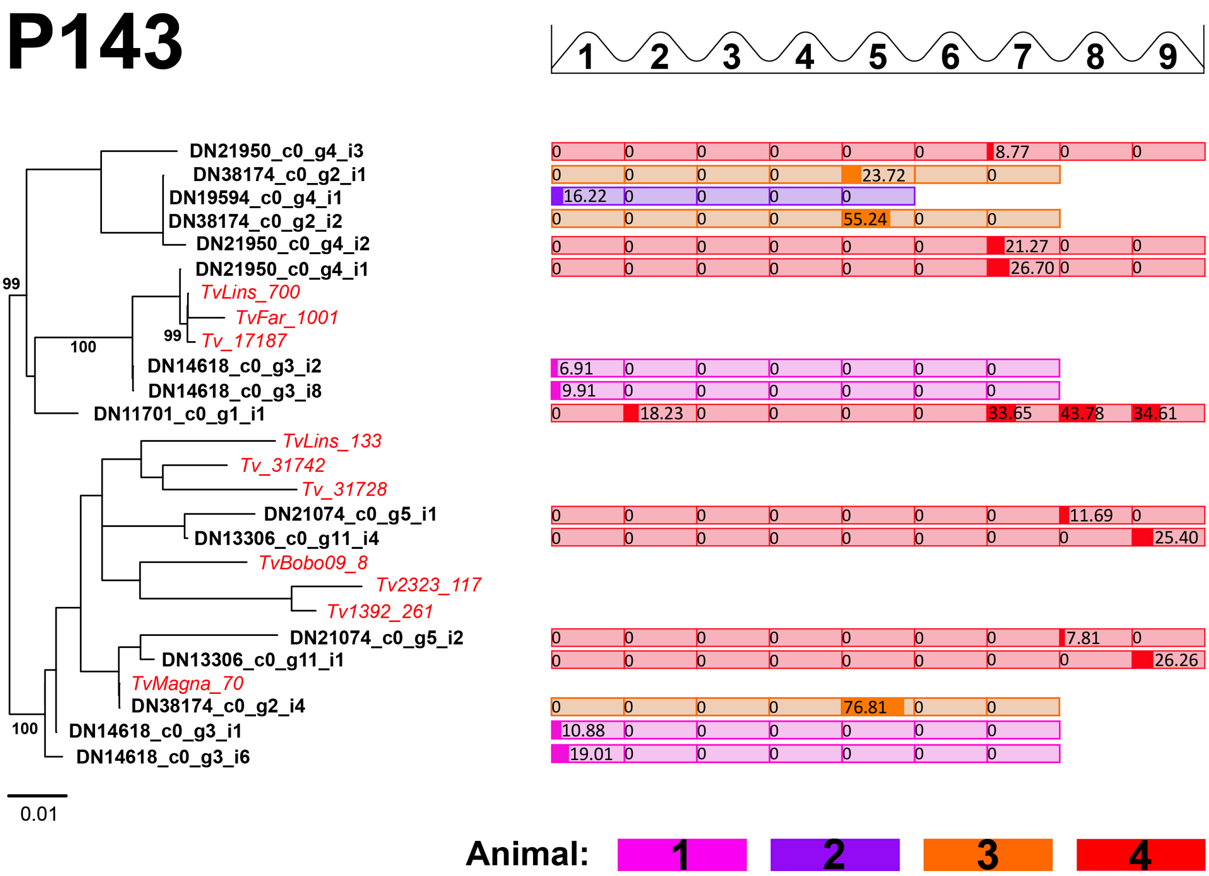


**Supplementary Fig. 5. Maximum likelihood phylogeny of Phylotype 143 showing the relationships among constituent genes from reference and strain genomes (COG type sequences, shown in red) and expressed *VSG* sequences from *T. vivax* Lins (shown in black).** The tree was estimated from a 618 bp alignment using a GTR+$\Gamma$ model in RAXML[2]. Robustness values (100 non-parametric bootstraps) are shown beside selected internal nodes. Beside terminal nodes representing expressed VSG are the transcript abundance values (CPM) for each peak of parasitaemia. These values are shaded by animal replicate.

P143 was observed at 7/29 peaks across four replicates and comprised an average of 2.71±1.3 transcripts per observation. Phylotypes were routinely observed to comprise multiple, distinct transcripts. The maximum number of unique P143 transcripts observed at a single peak was four in A1 (peak 1) and A4 (peak 9). P40 comprised a single transcript on 2/7 peaks when it was observed.

Phylotypes were observed to persist across consecutive peaks in the same animal but, typically, the actual transcripts concerned are distinct. However, P143 provides an unusual example of a single transcript re-emerging and persisting. DN11701 was expressed at peak 3 in A4 (one of several superabundant VSG), and then again, at a much lower level, at peaks 8, 9 and 10.

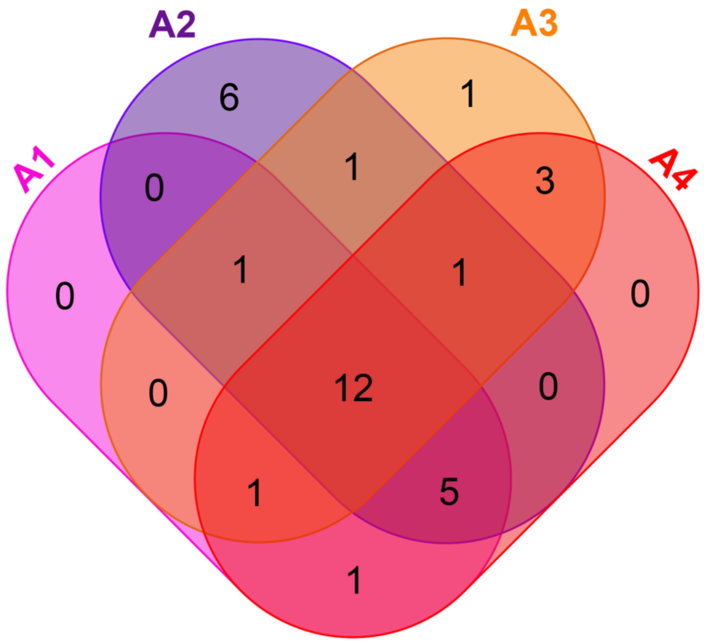


| Phylotype | A1 | A2 | A3 | A4 |
|---|---|---|---|---|
| 1 | 11 | 3 | 9 | 1 |
| 2 | 22 | 7 | 3 | 8 |
| 8 | 12 | 6 | 1 | 17 |
| 18 | 8 | 3 | 4 | 1 |
| 24 | 9 | 13 | 2 | 11 |
| 40 | 8 | 4 | 2 | 10 |
| 44 | 4 | 3 | 2 | 5 |
| 123 | 5 | 2 | 1 | 8 |
| 142 | 7 | 5 | 2 | 4 |
| 143 | 4 | 1 | 3 | 11 |
| 155 | 3 | 4 | 2 | 1 |
| 172 | 4 | 3 | 2 | 4 |
| 13 | 2 | 1 | - | 2 |
| 23 | - | 1 | 3 | 4 |
| 87 | 1 | 1 | - | 2 |
| 135 | 8 | 10 | 1 | - |
| 165 | 8 | 1 | - | 2 |
| 166 | 1 | 3 | - | 2 |
| 179 | 2 | 2 | - | 12 |
| 3 | 1 | - | 1 | 1 |
| 20 | 3 | - | - | 2 |
| 33 | - | - | 5 | 1 |
| 37 | - | - | 1 | 4 |
| 141 | - | 1 | 1 | - |
| 14 | - | 2 | - | - |
| 16 | - | 1 | - | - |
| 27 | - | 7 | - | - |
| 38 | - | 1 | - | - |
| 151 | - | 1 | - | - |
| 171 | - | 2 | - | - |
| 178 | - | - | 1 | - |

**Supplementary Fig. 6. A Venn diagram representing the phylotypes observed among expressed *VSG* transcripts in four animal replicates.** Individual phylotype distribution is tabulated alongside.

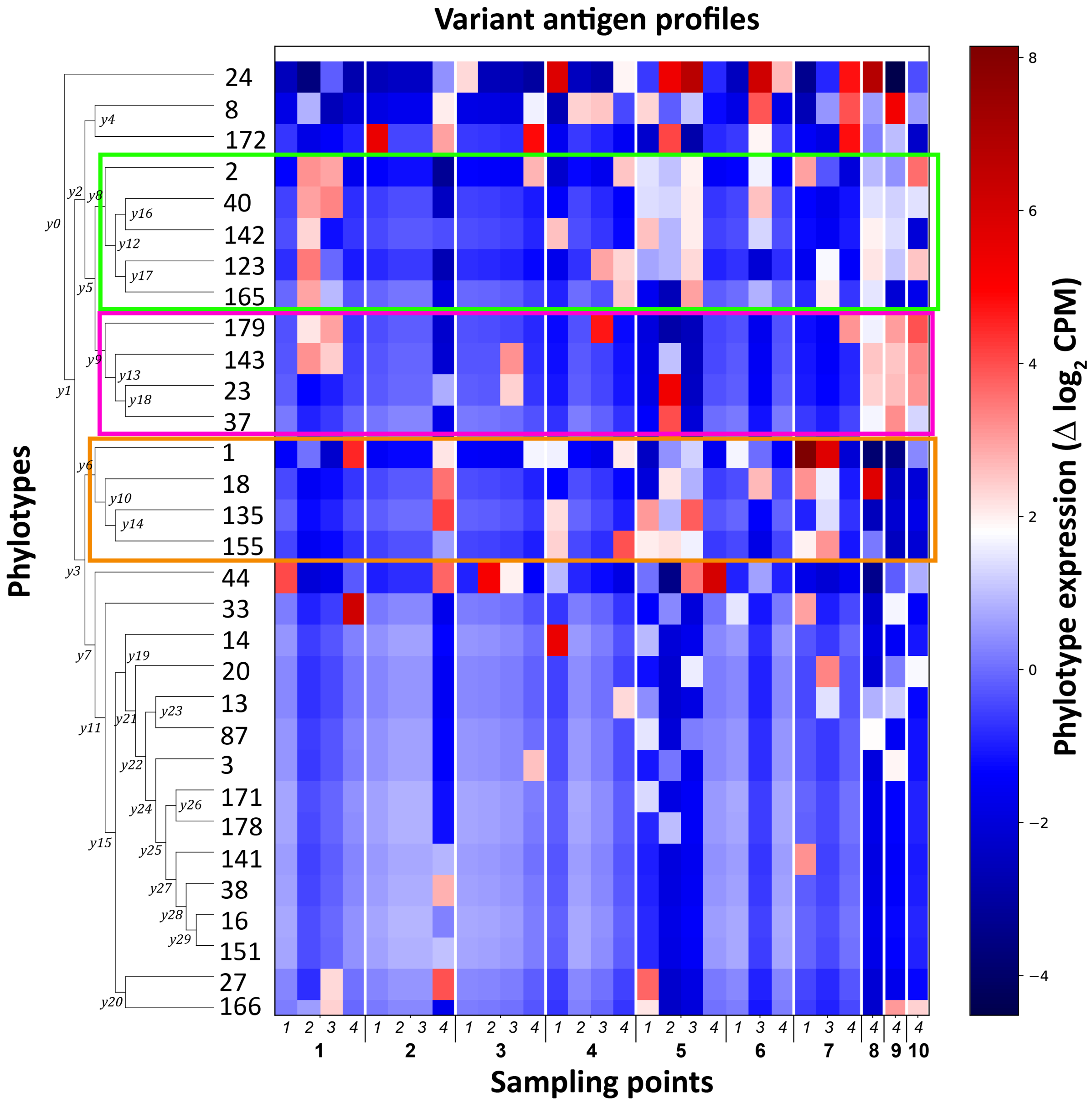


**Supplementary Fig. 7**. **A Gneiss (balance) analysis of phylotype expression across each animal**[3]**.** The heat map shows phylotype expression related to experimental time course. Cells in the map represent single sampling points from a given animal and peak of parasitaemia (x axis). On the x axis, animal number is shown in italics, and peaks of parasitaemia are numbered in bold. Cell shading represents changes in the combined transcript abundance from one peak to the next; red reflects increases in expression, blue represents decreases. Rows represent individual phylotypes; on the y-axis, these are arranged according to a dendrogram that clusters phylotypes based on Euclidean distances between phylotype profiles.

Each node of the dendrogram corresponds to a balance with each tip corresponding to a VSG phylotype. Significant differences in log ratio of time points were found in balances y1 (T8: β = -7.1, *p* = 0.04; T9: β = -6.6, *p* = 0.05) and y3 (T5: β = -3.4, p = 0.05; T7: β = -4.8, *p* = 0.01; T8: β = -6.3, *p* = 0.03), indicating that the repertoire of expressed phylotypes changed significantly at these points. However, there were no significant differences in balances y7, y11 and y15 suggesting that the log abundances of phylotypes expressed then were not significantly different between time points.

The figure shows that there was a modest reproducibility across animals in the identity and order of phylotypes, but substantial variation among replicates in the timing of expression. Bounded by a green line are examples of phylotypes (P2, P40, P142 and P143) expressed early (i.e. peak 1/2) in A2 and A3, that appear later at in A1-3 (peak 5/6), and even later in A4. A different set of phylotypes (P1, P18, P135 and P155) bounded by an orange line were expressed late (peak 5/7) in A1-3, but earlier in A4 (peak 2). In A4, where uniquely we have peaks 8-10, another set of phylotypes bounded by a pink line was expressed late (P179, P143, P23). Comparing these patterns to Fig. 5, it is notable that the phylotypes within these co-expressed groupings are often closely related.

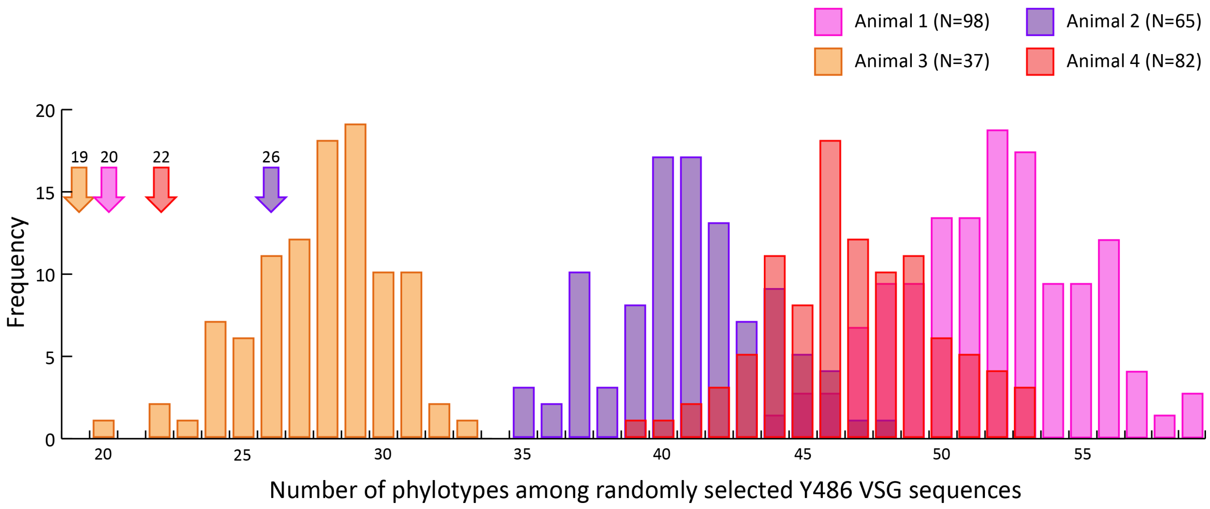


**Supplementary Fig. 8**. **Frequency distributions of the number of VSG phylotypes observed in simulated VSG repertoires.** VSG were selected at random from the Y486 repertoire to match the size of the observed repertoire and assigned to their phylotypes. This was repeated for each of four animal replicates. The observed number of phylotypes for each animal is indicated by arrows and is smaller than the simulated repertoires in each case. Furthermore, in each case the observed number lies outside of two standard deviations from the mean of the simulated repertoires. A1 expressed 20 phylotypes among 98 transcripts, simulated repertoires of the same size included mean average of 51.8 phylotypes (2xSD lower bound = 45.4). A2 expressed 26 phylotypes among 65 transcripts, simulated repertoires of the same size included mean average of 40.1 phylotypes (2xSD lower bound = 35.3). A3 expressed 19 phylotypes among 37 transcripts, simulated repertoires of the same size included mean average of 27.7 phylotypes (2xSD lower bound = 22.9). A4 expressed 22 phylotypes among 82 transcripts, simulated repertoires of the same size included mean average of 46.8 phylotypes (2xSD lower bound = 40.9).

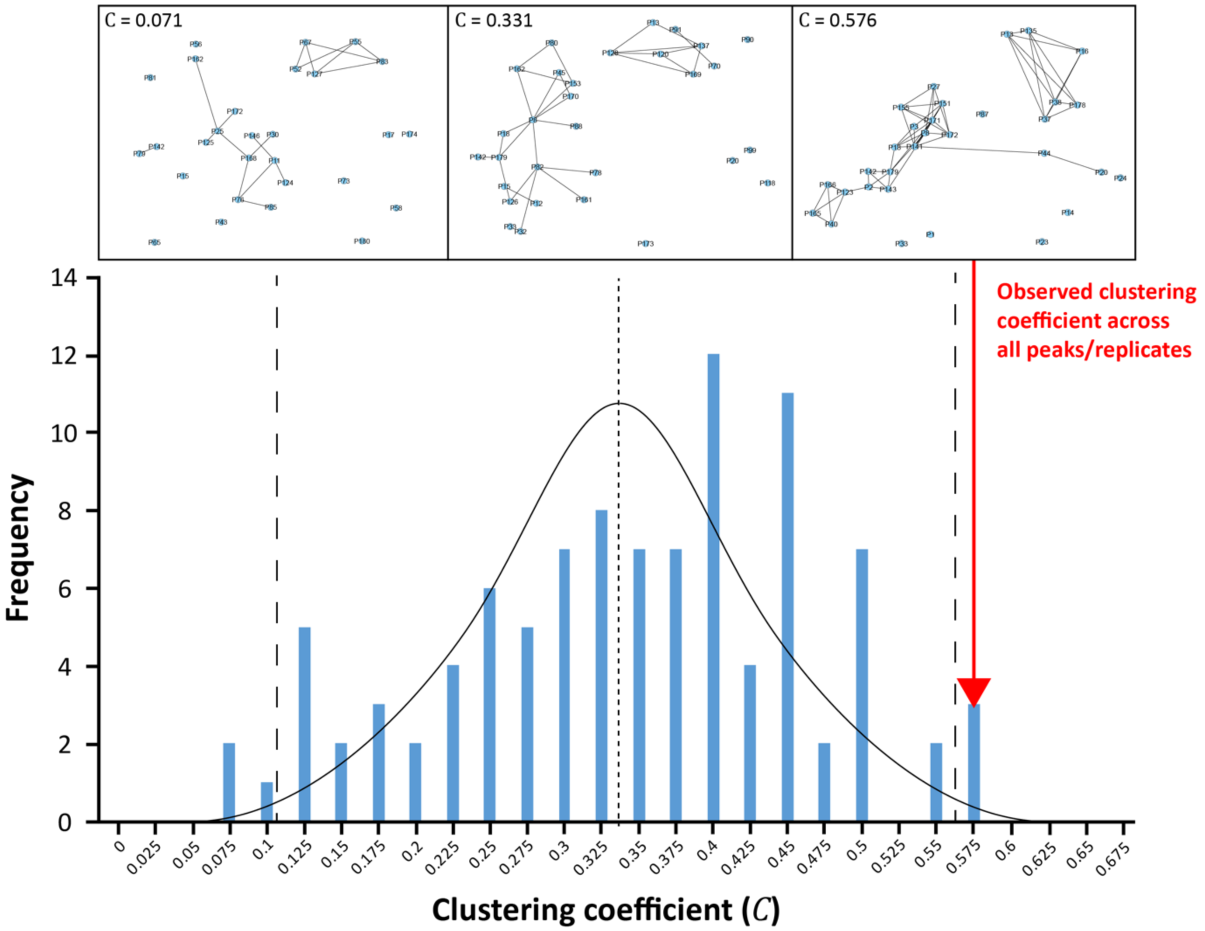


**Supplementary Fig. 9**. **Frequency distribution of clustering coefficients (C) for randomised sub-networks.** The phylotype networks shown in Figure 5 were analysed to compare the connectivity among 'expressed' nodes (i.e. those representing phylotypes that were expressed during experimental infections) with the connectivity expected by chance. C is a measure of the degree to which nodes in a graph tend to cluster together. The Network Analysis Tool in Cytoscape 3.7 was used to calculate the network average value for C, over all local clustering coefficients of all the vertices. Local clustering coefficients are computed as the proportion of connections among the immediate neighbours of a node that are realised compared with the number of all possible connections. Therefore, C is high where the nodes connected to a given node are themselves connected to each other. C was calculated for a subnetwork comprising all observed expressed nodes (N=31; red arrow). C was then calculated for 100 subnetworks of the same size where the nodes were selected by random number generator. The figure shows the distribution of C, which is approximately normal. Small and large dashed vertical lines represent the mean (0.333) and 2x standard deviations of the distribution respectively. The position of the observed value for C (0.576) exceeds twice the standard deviation of expected distribution (0.565), showing that the expressed nodes had a significantly higher value for C than would be expected by chance at $P < 0.05$. To illustrate, three subnetworks are shown above the distribution; a low-connectivity random subnetwork (left), a moderate-connectivity random network (centre) and the observed expressed node subnetwork (right).
